# 3D printed cell culture grid holders for improved cellular specimen preparation in cryo-electron microscopy

**DOI:** 10.1101/2020.06.12.147678

**Authors:** Florian Fäßler, Bettina Zens, Robert Hauschild, Florian K.M. Schur

## Abstract

Cryo-electron microscopy (cryo-EM) of cellular specimens provides insights into biological processes and structures within a native context. However, a major challenge still lies in the efficient and reproducible preparation of adherent cells for subsequent cryo-EM analysis. This is due to the sensitivity of many cellular specimens to the varying seeding and culturing conditions required for EM experiments, the often limited amount of cellular material and also the fragility of EM grids and their substrate. Here, we present low-cost and reusable 3D printed grid holders, designed to improve specimen preparation when culturing challenging cellular samples directly on grids. The described grid holders increase cell culture reproducibility and throughput, and reduce the resources required for cell culturing. We show that grid holders can be integrated into various cryo-EM workflows, including micro-patterning approaches to control cell seeding on grids, and for generating samples for cryo-focused ion beam milling and cryo-electron tomography experiments. Their adaptable design allows for the generation of specialized grid holders customized to a large variety of applications.

## Introduction

Cryo-electron microscopy (cryo-EM) and tomography (cryo-ET) allows for observing molecules or higher-order biological assemblies in their native, unperturbed state. In particular, cryo-focused ion beam milling scanning electron microscopy (cryo-FIBSEM) combined with cryo-ET shows great promise for revealing new biological structures and mechanisms within cells or even tissues (Rigort and Plitzko, 2015; Villa et al., 2013). While significant developments in cryo-FIBSEM/ET data acquisition and image processing have been made (Buckley et al., 2020; Hagen et al., 2017; Schur, 2019; Wagner et al., 2020; Zachs et al., 2020), sample preparation for complex cellular samples still often represents a major challenge. This is due to the sensitivity of many specimens to the varying seeding and culturing conditions required for electron microscopy experiments, as well as due to the difficulty of handling fragile, cell-compatible (i.e. not cytotoxic) EM grids. Cell types often studied in cryo-ET include small unicellular organisms such as *Chlamydomonas, Dictyostelia*, yeast or bacteria (for examples see (Engel et al., 2015; Jasnin et al., 2019; Kaplan et al., 2019). In addition, adherent mammalian cell lines are common specimens for ultrastructural analysis to understand cellular organization and mechanisms, such as organelle structure, intracellular transport or cell motility, to name a few (Azubel et al., 2019; Mahamid et al., 2016; Vinzenz et al., 2012). While non-adherent cells can be applied to EM grids immediately prior to vitrification using standard blotting approaches, more extensive cell culturing conditions are required for adherent cell monolayers that need to settle and attach to the substrate on EM grids. These conditions potentially include repeated media exchanges, application of selected effector molecules or drugs, and examination via light microscopy, which can introduce unwanted mechanical manipulations (such as repeated movement of cell culture dishes or displacement and relocation of grids). This can result in suboptimal culturing conditions and furthermore lead to distortion, bending or complete loss of grids. These considerations highlight the challenging nature of short and in particular long-term on-grid experiments using adherent cells.

In standard workflows EM grids commonly need to be placed in sufficiently large cell culture dishes, such as 6-well or 12-well plastic plates to ensure accessibility of grids for proper handling and retrieval. An additional layer of flexible film composed of waxes and polyolefins (i.e. Parafilm) can be used in cell culture dishes to provide an attachment surface for grids, to prevent flotation during seeding (Resch et al., 2011). This additional layer, however, often results in reduced visibility of cells, when seeding success is assessed via light microscopy. Moreover, due to the common use of excessively large cell culture dishes, the majority of seeding surface is not the desired target for cell adhesion (for example the surface area of commonly used 6-well plates is 9.6cm^2^ compared to 7.3mm^2^ of a single grid) and requires excessive quantities of media, cells and possibly effector molecules. This can become especially problematic if cells or effectors are not available in large excess, such as cells obtained from tissues, that are not commercially available (such as dendritic cells (Leithner et al., 2016)) or small molecules.

The excessively large surface area also poses an increased risk for undesired grid movement. The requirement of repeated physical manipulation of grids (and cell culture dishes) during various handling steps in cell culture experiments can result in the above mentioned detachment of grids from cell culture dish surfaces during media exchange or in the bending of grids and rupture of the grid support film. Hence, a significant level of user experience and a *steady hand* are required to achieve sufficient throughput, an appropriate cell density and to ensure minimal damage to the grids during specimen preparation.

A notable improvement for controllable cell seeding conditions on grids was recently introduced by EM grid compatible substrate micropatterning approaches (Engel et al., 2019; Toro-nahuelpan et al., 2019). Here, only defined areas on grids are made adherent or are specifically functionalized with extracellular matrix (ECM) proteins, allowing one to precisely control the positioning, shape, and size of adherent cells. The steps involved in micropatterning, however, lead to even more extensive grid handling, again increasing the risk of grid distortion and hence reduced throughput.

In the last decade, 3D printing of custom-designed tools has significantly augmented scientific workflows in a wide range of applications (Au et al., 2015; Huber et al., 2016; Jeandupeux et al., 2015). 3D printing using glycol-modified polyethylene terephthalate (PETG), which is also used for the manufacturing of cell culture vessels and other bio-compatible materials, allows its implementation in experiments including mammalian cells, where cytotoxicity or other altering effects need to be avoided. Further, cultured neurons exhibited no adverse effects over the course of several days when co-cultured with 3D printed poly-lactic acid (PLA), another commonly used material for 3D printing, (Gulyas et al., 2018).

Here, we introduce the application of low-cost, reusable 3D printed grid holders of varying shapes and dimensions for use in (cryo-)EM specimen preparation of cultured cells. The holders decrease the number of direct manipulations of grids during cell culturing experiments, allow for improved imaging of grids in culture using phase contrast and fluorescence imaging and enable the use of small volume cell culture dishes. As a result, this drastically reduces the required resources per grid. Here, we demonstrate that our cell culture grid holders integrate seamlessly into cryo-FIB and cryo-ET sample preparation workflows for different adherent cell lines. Moreover, they are compatible with micropatterning approaches for controlled cell seeding.

In summary, the application of grid holders facilitates sample preparation of sensitive cellular specimens for cryo-EM.

## Results

### Grid holder design

Experimental steps required for the preparation of adherent cells for electron microscopy include (Resch et al., 2011): **1)** Grid surface hydrophilization via glow discharging; **2)** Grid surface functionalization using ECM proteins (a step that can be omitted dependent on cell type); **3)** Placement of grids in cell culture dishes, usually on Parafilm as adhesive substrate; **4)** Cell seeding and (optional) treatment or washing; **5)** repeated light microscopic inspection of grids to check seeding conditions/success; and **6)** vitrification. In commonly employed workflows, these steps require repeated relocation of grids (e.g. from a plasma cleaner to cell culture dishes) and risk the danger of undesired grid movement (i.e. detachment of the grid from Parafilm when adding cell culture medium).

When designing our grid-holders for cell culture experiments we aimed for an easily adaptable design that minimizes direct grid handling steps, to placing grids into, and retrieving them from holders, as the two only direct handling steps. At the same time, we intended to reduce the required number of cells and the quantity of reagents. In addition, we wanted to provide users with the opportunity to perform both short, and also long-term culturing experiments (up to weeks if necessary) using the described holders, hence facilitating repeated specimen treatment (e.g. with transfection reagents) or washing and imaging of seeded cells. Finally, we intended to make our grid holders compatible with ‘on-grid’ micropatterning approaches to ensure optimal seeding and spreading of adherent cells on grids. While these prerequisites put several limitations on the overall grid holder dimensions and also required defined topological features for easy placement and retrieval of grids, the designs shown are but examples for an essentially limitless number of designs for different experimental setups.

Specifically, two different designs of cell culture compatible grid holders are shown in Figure 1, which can be routinely used in various experimental workflows. Both designs feature a recess into which a single EM grid can be stably placed, substantially reducing any additional grid movement throughout subsequent handling steps. An opening at the base of the grid holders allows imaging the grid center (the region usually desired for follow-up experiments in cryo-ET) on any cell culture microscope in phase contrast and fluorescence settings (Figure 1A,B). The grid holders are specifically tailored to fit 24- or 96-well plates, occupying the majority of space within the wells, additionally limiting the freedom of movement of the holders themselves and significantly reducing the required volumes of media and cells (Figure 1C). Both PLA and PETG have a density of 1.24g/cm^3^ and 1.38g/cm^3^, respectively, and hence holders do not float in cell culture dishes when printed with sufficiently high infill percentages. For the circle-base grid holders (Figure 1, left panel) an additional extension on the top is added for facilitating retrieval from narrow 96-well plates. The grid holders can be produced in commercially available 3D printers with printing times of few minutes and at material costs of only cents per grid holder.

**Figure 1:**
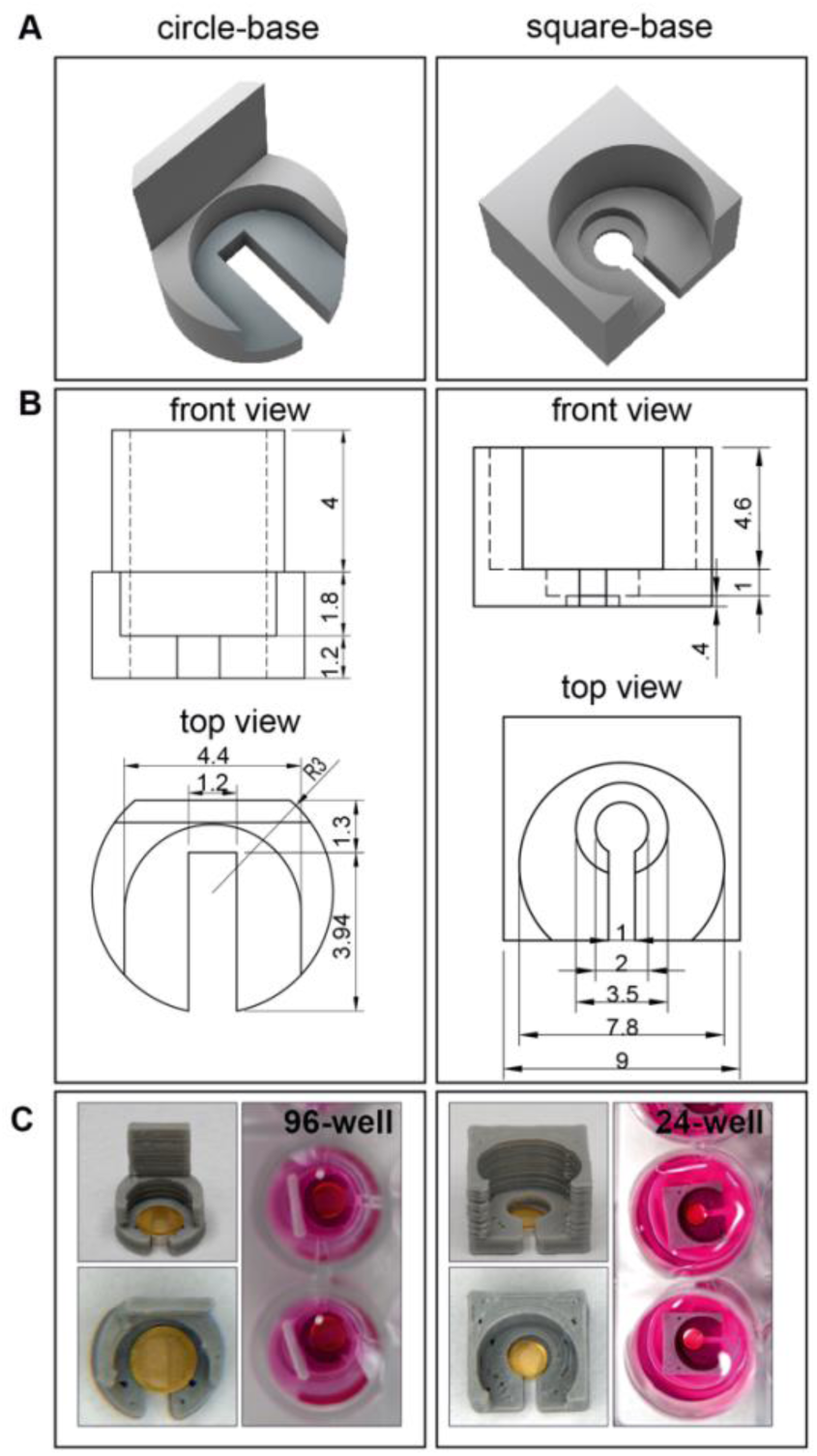
Design of 3D printed grid holders. **A)** Schematic representation of two different grid holder designs. **B)** Technical drawings of the shown designs. Dimensions are indicated in the front and top views and are given in millimetres. Note the difference in size between the circle and square grid holders. **C)** Photographs of grid holders with loaded grids in standard 96- and 24-well cell culture plates.

### On-grid culturing using grid holders

We subjected our grid holders to a complete experimental workflow using HeLa cells to compare them to a commonly used approach (Resch et al., 2011), in which gold grids are placed in 6-well plates on Parafilm (Figure 2A). Upon comparison of imaging conditions in phase contrast or fluorescence microscopy, grid holders offer improved visibility of cells, such as enhanced distinguishability of cells sitting on grids or below on the well plate surface (Figure 2A, left and middle panel). Additionally, grids derived from our grid-holder workflow consistently showed a high level of integrity (judged from the completeness of the EM grid support film using TEM analysis) and cells were distributed evenly over the grid (Figure 2A, right panel).

**Figure 2:**
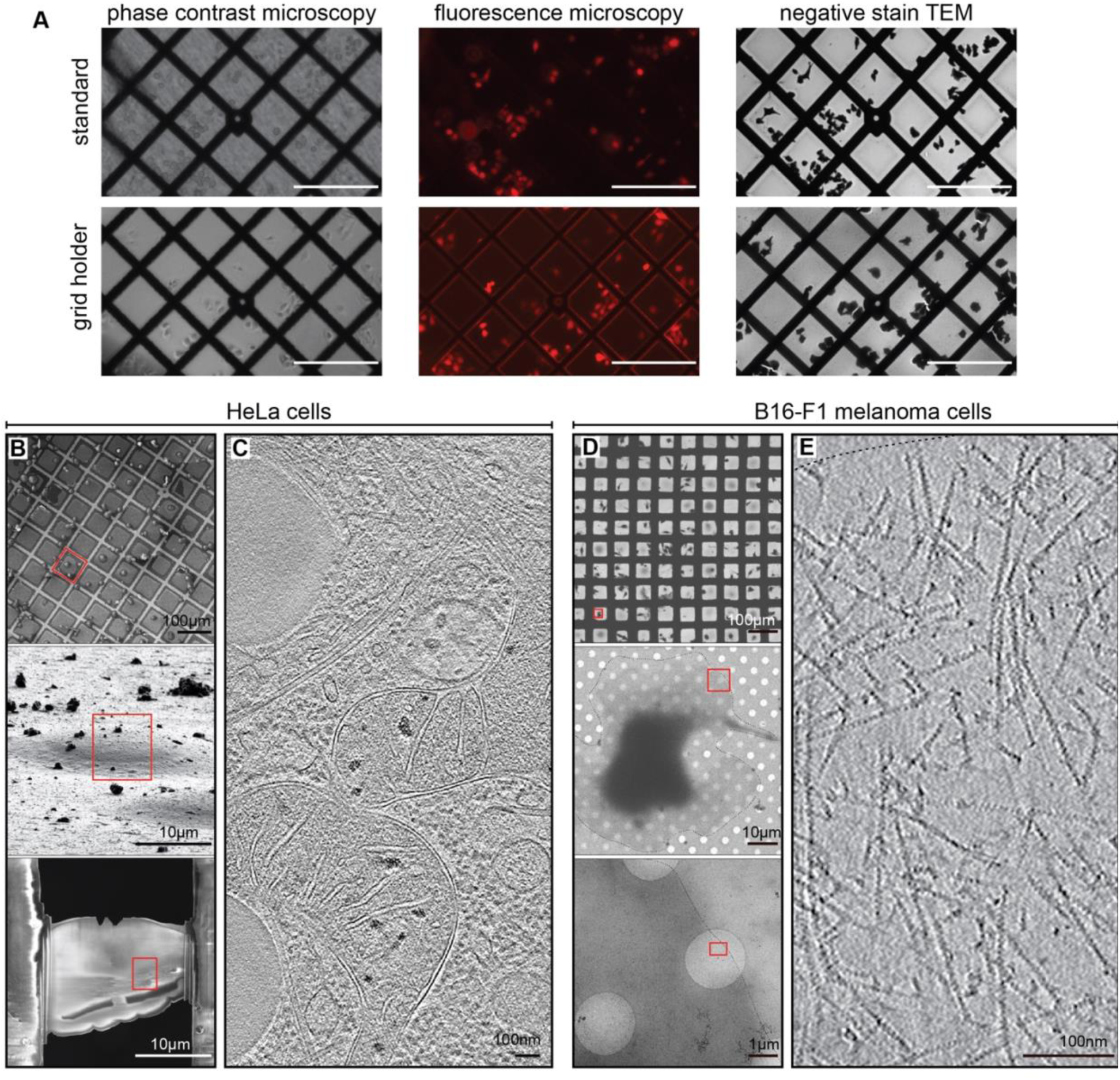
Compatibility of grid holders with standard cryo-EM workflows. **A)** Comparison of cell seeding and culturing of mCherry-expressing HeLa cells on Formvar coated grids using a standard workflow on parafilm (top) and our grid holders (bottom). Visibility of cells on grids in grid holders is improved in phase contrast microscopy (left) and fluorescence microscopy (middle). The integrity of grids and support film is further shown via negative staining TEM (right). Scale bars, 100µm. **B-C)** cryo-FIBSEM and cryo-ET of HeLa cells prepared using a square-based grid holder. **B)** Cryo-SEM images of Hela cells shown at increasing magnifications. Lower panel in **B** displays the finished lamella (∼200nm thickness) prepared from the cell shown in the middle panel. **C)** Sum of 7 computational slices through a tomogram of the lamella depicted in **B** and showing the organization of the nuclear periphery. **D-E**) cryo-ET of B16 cells. **D)** cryo-TEM of B16 cells shown at increasing magnifications, from a low-magnification overview to a zoom in of the leading edge of a selected cell. **E)** Sum of 5 computational slices through gaussian-filtered tomogram of the specified area in the lowest panel of **D**, showing the organization of branched actin filaments within a lamellipodium. **B-E)** Scale bars sizes are annotated.

One main objective for the design of the grid holders was the reduction of the total number of cells and amount of medium and reagents needed, when compared to a standard workflow. Using grid holders, we achieved a similar seeding density using an order of magnitude fewer cells (i.e. tens of thousands of cells versus only few thousands in the standard and grid holder workflow, respectively, see *Materials and Methods*). Additionally, use of cell culture reagents could be reduced substantially, such as from 2mL of medium per sample in a 6-well plate to 100µL in a 96-well plate. Moreover, we confirmed in 2 independent repetitions that even when incubating cells in a small volume of 100µL, cell proliferation was unaffected by the presence of PETG and PLA material (one representative repetition: 3.36 proliferations per day in control cells vs. 3.75 in cells incubated with PLA and 3.26 with PETG; one-way-ANOVA, n=10, F=0.3507 and p=0.7073).

### Ultrastructural and morphological characterization

We then examined cell morphology and ultrastructural preservation of HeLa cells and B16-F1 melanoma cells (a routinely used cell model to study cell migration (Damiano-Guercio et al., 2020; Dimchev et al., 2020; Koestler et al., 2008)) cultured on grids using the described holders by cryo-EM. As expected, the use of grid holders did not affect cell morphology. HeLa cells robustly settled onto grids and could be subjected to subsequent FIB-milling and cryo-ET analysis (Figure 2B,C). B16-F1 cells, which form thin cellular protrusions called lamellipodia at their front, spread out evenly on the grid (Figure 2D). To visualize subcellular structure at the leading edge of these B16-F1 cells we used a protocol for extraction and fixing that optimally retains the actin network for ultrastructural analysis (Koestler et al., 2008, see *Materials and Methods* for details). Zooming into the leading edge of cells allowed the clear visualization of actin filaments within lamellipodia using cryo-ET (Figure 2E), similar to previous descriptions of adherent cells (Vinzenz et al., 2012).

For both experiments grid holders resulted in a facilitated and streamlined sample preparation process, with a high number of intact samples due to reduced direct grid handling steps, and without causing any disadvantages to cell morphology or viability.

### Micropatterning-compatible grid holders

Since EM grid micropatterning shows great promise for further simplified and more efficient cell seeding (Engel et al., 2019; Toro-nahuelpan et al., 2019) we designed grid holders that are also compatible with micropatterning approaches using a standard, non-commercial setup. Specifically, high-throughput patterning on EM grids requires grid holders to provide a low, stable and even support of grids for controlled ablation of the Polyethylene glycol(PEG)-brush layer, which is applied to EM grids to render them non-adherent. Hence, we re-designed the square grid holder shown in Figure 1A to have a base height of only 0.2mm allowing positioning of grids as close as possible to the bottom of optical well plates. This ensured accurate focusing of the pulsed laser illumination on the PEG layer and its ablation. Using our micropatterning setup we were then able to create patterns of different shapes and dimensions (Figure 3A, left panels). Using our customized pattering setup, we achieved a patterning rate of 1000µm^2^/50s, resulting in roughly 6 minutes ablation time per grid with 16×400µm^2^ square patterns.

**Figure 3:**
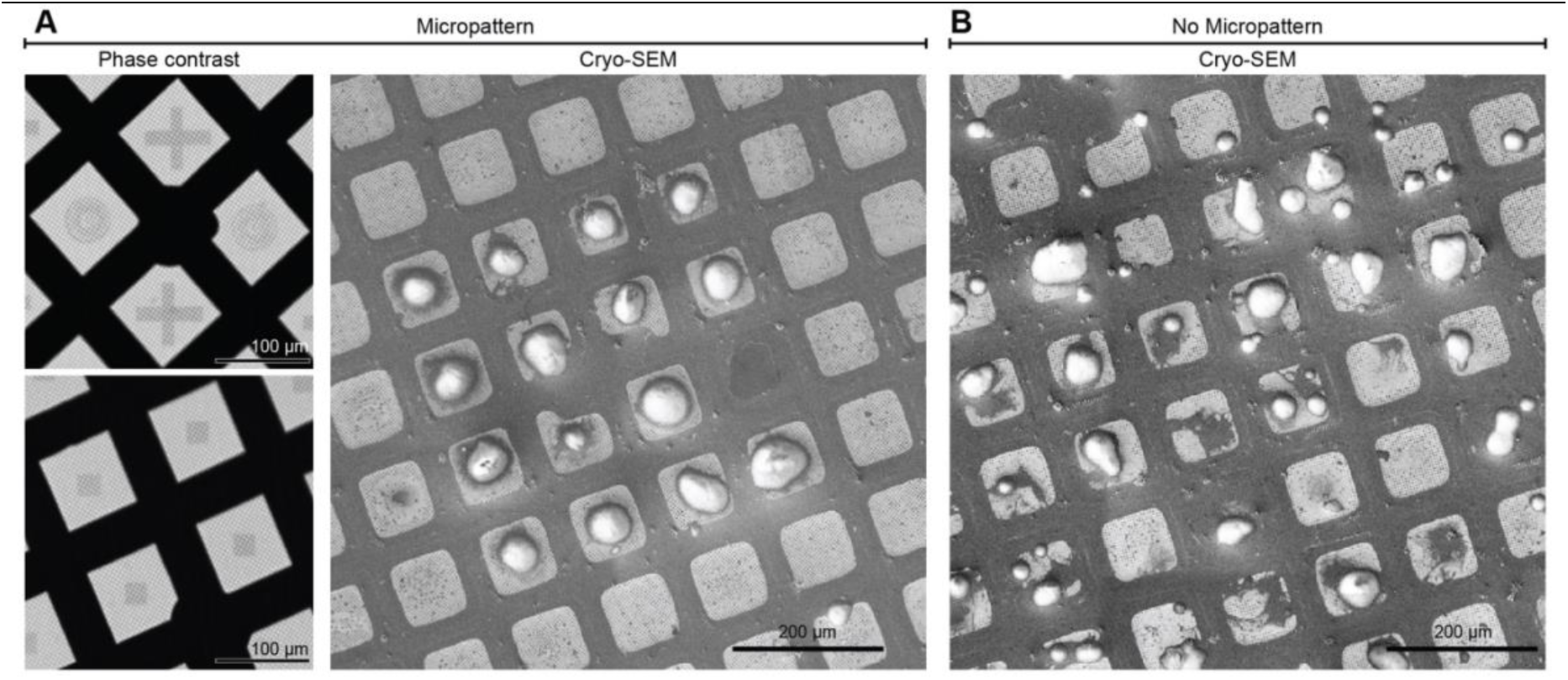
Micropatterning using grid holders. **A)** Patterns of different shapes and sizes ablated onto PEG-coated titanium SiO_2_ grids as seen via phase contrast light microscopy (left). NIH3T3 fibroblasts seeded onto square patterns additionally functionalized with fibronectin, visualized via cryo-SEM. The cells adhere only onto the patterned areas. **B)** Cell seeding of NIH3T3 fibroblasts on a non-patterned grid displays irregular seeding with many cells sitting close to the grid bar. Scale bar sizes are annotated in the figure.

Use of grid holders resulted in high patterning throughput, since moving between grids, which could be kept in adjacent and regularly spaced wells of one optical plate, was possible within seconds without requiring significant readjustments upon relocation. Additionally, using grids holders within 96-well plates reduced the risk of grids drying out during patterning, as all grids could be suspended in sufficiently large volumes. We then functionalized the patterned grids with fibronectin and seeded NIH3T3 fibroblast cells on top of these grids. Successful micropatterning and cell seeding was verified using phase-contrast microscopy and cryo-SEM, respectively (Figure 3A,B).

Again, the number of direct grid handling steps was reduced with the use of grid holders, limiting the risk of damage to grid or support film during sample preparation before, during and after patterning.

## Discussion

Here we introduce grid holders as tools for an easier and more resource-efficient preparation of cellular specimens for (cryo)-EM experiments. We show that our grid holders are compatible with different cell culturing workflows and increase throughput as well as reproducibility. Firstly, grid holders effectively reduce the number of direct grid handling steps and increase ease of media exchange or washing steps. Secondly, grid holders require only a fraction of cells, media and effectors/inhibitors compared to established workflows (Resch et al., 2011). Thirdly, the presented grid holders restrict movement and, due to the absence of Parafilm, allow for facilitated light microscopy observation of grids. They enable researchers to quickly, safely and unambiguously assess (and adapt) cell density during and after seeding on a single grid level. This represents a considerable advantage, as the risk of damaging grids and their support film during assessment of seeding conditions and culture maintenance during long-term experiments is mitigated. We also show the applicability of grid holders to micropatterning workflows (Figure 3), in which they allow automation of patterning on regularly spaced grids to additionally increase throughput.

While we describe the use of grid holders in the context of specimen preparation for cryo-ET, these devices are also applicable to any other workflow preceding room-temperature EM experiments, making them compatible with ACLAR® foil, sapphire discs or in theory even glass cover slips. Another advantage of the grid holders is that they can be introduced into the workflow at any given point, such as the start of an experiment where grids can be glow discharged within grid holders, or at a later step, such as after the application of ECM proteins or the seeding of cells. This makes them highly adaptable to the needs of the user for a wide range of applications.

Due to the availability of inexpensive 3D printers and open source/free academic license 3D design tools (for examples used here see *Material and Methods*), grid holders can be easily adapted for other experimental setups. This would for example allow culturing of different cell (geno-)types within the same well on different grids (Figure 4), ensuring equal experimental treatment of knock-out and wild type specimens. Other grid holders could be shaped to allow for seeding of two different cell types on the same grid, leading to simultaneous analysis of both samples.

**Figure 4:**
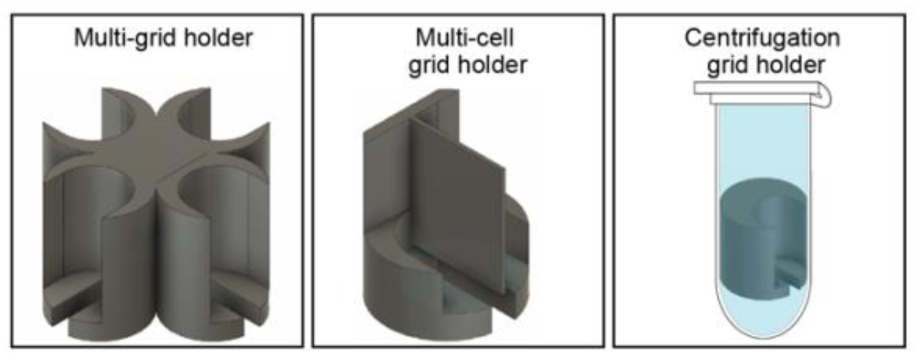
Versatility of designs of 3D printed grid holders. A selection of different grid holder designs is shown, allowing various experimental setups.

For other applications (i.e. studying non-adherent cells), holders can be tailored to fit into 1.5 or 2ml reaction tubes allowing for fast sedimentation of samples via centrifugation. Incubation could then be performed within the tube or after transfer via the grid holder in a well plate. Furthermore, grid holders could be adapted to fit into microfluidic devices, allowing for analysis of cells on grids during chemotaxis.

3D printing has a profound impact in everyday life and science, with continuous and rapid development of new designs (Ngo et al., 2018). Similar to other recent developments for controlled cell seeding onto EM grids, we therefore expect that the grid holders introduced here can become a standard tool in the preparation of cellular specimens as they harbour the potential to facilitate grid handling, increase sample quality and enable novel experimental designs. Development of highly specialized grid holders will be a matter of time and are supported by the availability of open source/free 3D design software and affordable printers.

## Acknowledgements

Bettina Zens acknowledges support by the Niederösterreich Fond. This research was supported by the Scientific Service Units (SSU) of IST Austria through resources provided by Scientific Computing (SciComp), the Life Science Facility (LSF), the BioImaging Facility (BIF) and the Electron Microscopy Facility (EMF). We thank Georgi Dimchev and Sonja Jacob (Vienna Biocenter Core Facilities) for testing our grid holders in different experimental setups and Daniel Gütl and the Kondrashov group (IST Austria) for granting us repeated access to their 3D printers. We are thankful to Ori Avinoam and William Wan for helpful comments on the manuscript and also thank Dorotea Fracchiolla (Art&Science) for illustrating the graphical abstract.

## Author Contributions

FF, BZ and FKMS conceived the project and designed the experiments. FF and BZ designed grid holders, performed cell culture and cryo-EM experiments and analysed the data, with input of FKMS. RH and BZ established the on-grid micropatterning experiments. FF, BZ and FKMS wrote the manuscript.

## Materials and Methods

### Design and 3D-printing of grid holders

Grid holders were designed in Autodesk Fusion 360 (free education license) and printed with either an Original Prusa MINI (Prusa Research) or BLV mgn Cube (open source 3D printer project). Individual grid holders were printed in a matter of minutes using 100% infill. For printing, either PLA (Prusa Research) or PETG (Filament PM) filaments were used. Grid holders were sterilized with perform® classic alcohol EP and UV irradiation prior to use in cell culture experiments. Grid holders have been re-used up to 15 times without any signs of negative effects on sample preparation.

Plans for the presented grid holders are available online at https://schurlab.ist.ac.at/downloads with a creative commons CC BY-NC-SA 4.0 license.

### Cell Culture and cell seeding

Wildtype B16-F1 melanoma and wildtype NIH 3T3 cells, as well as HeLa cells expressing cytosolic mCherry were kindly provided by Klemens Rottner (Technical University Braunschweig, Helmholtz Centre for Infection Research) and Michael Sixt (IST Austria), respectively. Cells were cultured in Dulbecco’s modified Eagle’s medium (DMEM GlutaMAX, ThermoFischer Scientific, #31966047), supplemented with 10% (v/v) fetal bovine serum (ThermoFischer Scientific, #10270106) and 1% (v/v) penicillin-streptomycin (ThermoFischer Scientific, #15070063). Cells were incubated at 37°C and 5% CO2.

Either 200 mesh gold holey carbon grids (R2/2; Quantifoil Micro Tools) or 200 mesh gold grids (Science Services, #G200-AU) coated with continuous Formvar film (0.75%) were used. Throughout all cell culture experiments Dumont tweezers, medical grade, style 5 and style 7 were used.

Prior to seeding of cells, grids were glow discharged in an ELMO glow discharge unit (Cordouan Technologies) for 2min (holey carbon), or 30s (formvar), either directly in the specific grid holders or on Parafilm. Grids were then coated with 25µg/ml fibronectin (Sigma-Aldrich, #11051407001) for 1h at 37°C.

Circle-base grid holders with grids were then placed in 96-well plates and 80µl medium was added on top of the grids. A total number of 2.5×10^3^ cells were seeded onto the grid. Square-base grid holders with grids were placed into 24-well plates containing 800µl medium per well and then seeded with a total number of 1.5*10^4^ cells. Cells were allowed to settle for 3h prior to imaging and further sample preparation. Cell viability and seeding was observed via the EVOS cell imaging system (Evos® fl, peqlab). For cell seeding and culturing without grid holder (i.e. a standard workflow), the bottom of 6-well plates was covered with parafilm according to a previously published protocol (Resch et al., 2011), and one grid was placed in each well. 3ml of a cell suspension containing a total of 2.5×10^5^ cells was added per well and mixed by gentle shaking. Cells were allowed to settle for 4h prior to imaging and subsequent sample preparation steps.

### Viability Test

30 wells of a 96-well plate were seeded with 2.5×10^3^ cells per well and cells were left to settle for 4h prior to imaging at time point 0. Subsequently, 10 rings printed from PLA were placed in a randomized fashion into single wells before 10 rings made of PETG were placed in 10 other randomly chosen wells. Rings were designed to have a comparable surface to circle-base grid holders. 10 wells were incubated without 3D printed material as control. After 24h, each well was imaged again. Cells were counted with ImageJ (Schneider et al., 2012) using the Multi-Point tool and statistical analysis was performed in GraphPad Prism. Data was tested for normal distribution with a normality test, and for variance with a Browne-Forsythe test. Subsequently, the sample groups of control, PETG and PLA incubation were subjected to an ANOVA test, which revealed no significance. The experiment was performed in 2 independent repetitions, yielding comparable results.

### Micropatterning

200 mesh holey silicon dioxide (SiO_2_) titanium grids (R2/2; Quantifoil Micro Tools) were glow discharged in an ELMO glow discharge unit (Cordouan Technologies) for 2min directly in micropatterning grid holders. Grids were then coated with Polyethylene glycol (PEG)-brush (SuSoS AG, #PLL(20)-g[3.5]-PEG(5)) at a concentration of 1mg/ml in PBS (Thermo Fischer Scientific, #20012019) for 1h at room temperature (RT) and subsequently dried briefly using filter paper. Grids were turned 180 degrees upside down in their grid holders to have the PEG coated side facing the optical bottom of a µ-96-well plate (ibidi, # 89626) for laser ablation. 40µl PBS were added to each well, to keep the grids from drying out during the ablation procedure.

The laser ablation setup consisted of a pulsed 355nm UV laser (PowerChip, Teem Photons) coupled to the rear port of an inverted microscope (Axio Observer Z1, Zeiss). The laser power was attenuated to an average power of 6 µW (with pulse frequency of 1kHz and pulse duration <350 ps) using an acousto-optic modulator (AA.MQl l0-43-UV, Pegasus Optik). A long working distance dry objective (LD Plan-Neofluar 20x/0.4, Zeiss) together with a pair of galvanometric mirrors (Lightning DS, Cambridge Technology) was used to focus the UV laser anywhere in the sample plane.

The ablation parameters were chosen to achieve a visible darkening of the ablation area, when observed in brightfield transmission, while staying well below the destruction threshold of the EM grid. For line features the laser was focused on the SiO2 layer for a duration of 10 pulses per shot and 2 shots per µm^2^. For areal structures the energy density was reduced by decreasing the shots per µm^2^ to 1.

After micropatterning, grids were again flipped in their grid holders, dried again briefly with filter paper and coated with 25µm/ml fibronectin for 1h at RT. Grids were then washed once with PBS and cell culture medium. Grids were incubated with 150µl of cell culture medium for 30min at 37°C prior to seeding of 1*10^4^ NIH 3T3 cells on top of the grids. Cells were allowed to settle for 3h, before phase contrast imaging to verify seeding success, and vitrification as described below.

### (Cryo)-electron microscopy

Negative staining of HeLa cells was performed by dropwise application and immediate blotting of 50µl total volume of 4% negative-staining-solution (10 nanometre BSA conjugated gold colloid diluted in 4% sodium silicotungstate, pH7.0, Agar Scientific, #AGR1230). Grids were subsequently imaged on a TEM Tecnai 10 operated at 80kV and equipped with a LaB6 filament and an OSIS Megaview III camera.

For cryo-EM analysis of B16-F1 melanoma cells (shown in Figure 2), cells were prepared as previously described (Koestler et al., 2008). In brief, grids were removed from the grid holders, placed in a 50µl drop of cytoskeleton buffer (10mM MES, 150mM NaCl, 5mM EGTA, 5mM glucose and 5mM MgCl_2_, pH6.2) with 0.75% Triton X-100 (Sigma-Aldrich, #T8787), 0.25% glutaraldehyde (Electron Microscopy Services, #E16220) and 0.1µg/ml phalloidin (Sigma-Aldrich, #P2141) and incubated for 1min at RT. Grids were post-fixed in a 50µl drop of cytoskeleton buffer containing 2% glutaraldehyde and 1µg/ml phalloidin for 15min at RT.

Grids containing unfixed HeLa or NIH3T3 cells and extracted/fixed B16-F1 melanoma cells were vitrified in liquid ethane after backside plotting in a Leica GP2 plunger (Leica Microsystems). Plotting conditions were set to 37°C, 90% humidity, 3.5s blot time (NIH3T3 or HeLa cells) or 4°C, 80% humidity, 2.5s blot time (B16-F1 melamona cells). 10 nanometre BSA conjugated gold colloid diluted in PBS was applied prior to blotting onto B16-F1 melamona cell samples to serve as fiducials during tomogram reconstruction. Vitrified grids were clipped into autogrids and stored under liquid nitrogen conditions until further processing.

For cryo-fluorescence imaging (to check integrity of grids and choose targets for ion beam milling), clipped grids were imaged on a Leica Cryo CLEM microscope (Leica Microsystems) using the Leica Application Suite 3.7.0. Lamellae were generated using a Thermo Scientific Aquilos cryo-DualBeam (focused ion beam/scanning electron microscope). Cryo-light and cryo-SEM images were correlated using the MAPS 3.3 software (Thermo Scientific).

For ion-beam milling, samples were first coated with organometallic platinum using the *in situ* gas injection system (15s at WD 2.5mm). Lamella were prepared using the Gallium ion beam at 30kV and a stage tilt angle of 17°, in a stepwise manner, starting with a current of 1nA and gradually reducing the current to 30pA for the final cleaning steps. Progress of the milling process was monitored using the SEM beam at 2keV and 25pA until a desired thickness of ∼200nm was obtained. Grids were then stored in liquid nitrogen conditions until further use for cryo-electron tomography.

Cryo-ET data on lamellae was collected on a Thermo Scientific Glacios TEM operated at 200kV and equipped with a Falcon 3 camera operated in linear mode using SerialEM (Mastronarde, 2005). Tilt series were acquired with a bidirectional scheme starting from 0º, with an increment of 3º and total range from −60º to +60º. The nominal magnification was 22,000x, resulting in a pixel size of 6.511Å. The total dose was ∼120e/Å^2^ and the nominal defocus was set to −8µm.

Cryo-electron tomograms on extracted and fixed B16 cells were acquired on a Thermo Scientific Titan Krios TEM equipped with a BioQuantum post column energy filter and a K3 camera (Gatan) using SerialEM (Mastronarde, 2005). The slit width of the filter was set to 20eV. Tilt series were acquired with a dose-symmetric tilt scheme ranging from −60 ºto 60º with a 2º increment (Hagen et al., 2017). The nominal magnification was 42,000x, resulting in a pixel size of 2.137Å. The cumulative dose was 180e/Å^2^ and the nominal defocus was set to −4μm. Individual tilt images were acquired as 11520×8184 super resolution movies of seven frames, aligned on-the-fly during data acquisition using the SerialEMCCD frame alignment plugin and saved as 2x binned mrc stacks.

Tomogram reconstruction and visualization was done using the IMOD software package (Kremer et al., 1996).

